# Transfer Learning from Markov models leads to efficient sampling of related systems

**DOI:** 10.1101/158592

**Authors:** Mohammad M. Sultan, Vijay S. Pande

## Abstract

We recently showed that the time-structure based independent component analysis method from Markov state model literature provided a set of variationally optimal slow collective variables for Metadynamics (tICA-Metadynamics). In this paper, we extend the methodology towards efficient sampling of related mutants by borrowing ideas from transfer learning methods in machine learning. Our method explicitly assumes that a similar set of slow modes and metastable states are found in both the wild type (base line) and its mutants. Under this assumption, we describe a few simple techniques using sequence mapping for transferring the slow modes and structural information contained in the wild type simulation to a mutant model for performing enhanced sampling. The resulting simulations can then be reweighted onto the full-phase space using Multi-state Bennett Acceptance Ratio, allowing for thermodynamic comparison against the wild type. We first benchmark our methodology by re-capturing alanine dipeptide dynamics across a range of different atomistic force fields, including the polarizable Amoeba force field, after learning a set of slow modes using Amber ff99sb-ILDN. We next extend the method by including structural data from the wild type simulation and apply the technique to recapturing the affects of the GTT mutation on the FIP35 WW domain.

## Introduction

Efficient sampling of protein configuration space remains an unsolved problem in computational biophysics. While algorithmic advances in molecular dynamics (MD) code bases^1^ combined with distributed computing hardware^2^, specialized chips^3^, and large-scale increasingly faster GPU clusters have provided routine access to microsecond timescale dynamics, there is still room for significant improvements.

One such potential avenue is predicting the effects of mutations onto the protein’s wild type free energy landscape. At this point it is worth explicitly noting that while we use the biological terms wild type and mutant extensively, our methodology is easily generalizable to scenarios where a baseline (aka the wild type) free energy landscape has been mapped out and we now wish to understand the dynamical consequences of a making some changes to system (aka the mutation). Under the current scheme, one would have to re-run our entire simulation to ascertain the affects of a mutation onto a protein’s free energy landscape. Due to the vast amount of computational resources required for even one simulation, most current MD papers run one simulation in a single force field for a single protein. However, considering the important role of mutagenesis experimentally as biophysical probes, the biological role of SNPs in medicine and disease, as well as phylogenic and evolutionary questions connecting mutations, often there are hundreds to thousands of mutations (or more) that would be relevant for simulation. Instead, the predictions from these simulations are extrapolated to other conditions but those changes/mutations are often not explicitly tested in-silico. While such hypothesis generation is useful for guiding future work, the gap between extrapolated predictions and experimental realization is large.

There is an obvious scaling problem between the computational and time cost of unbiased MD and the number of interesting mutants that could be investigated using simulation. For example, there are several hundred known protein kinases^4,5^ with each having tens to hundreds of known mutants. These kinases have critical protonation and phosphorylation sites that significantly affect their free energy landscapes^6,7^. To predict these mutations’ effects, do we need to re-run an entirely new simulation on the mutated protein? Are modern force fields even capable of elucidating such effects? Even if we assume an accurate enough force-field^8,9^, how do we efficiently sample these mutants or perhaps even propose new novel variants to be probed via experiments. Arguably, for MD to decrease the gap between theoretical hypothesis and experimental realization, an ability to efficiently sample the effects of mutations is required. Since unbiased MD is too slow, we turn to enhanced sampling.

While enhanced sampling methods such as Metadynamics or Umbrella sampling offer promise, they require identification of a set of collective variables (CVs)^10^ to sample along. Metadynamics^10–13^ can be thought of as computational sand filling along CV of interest to enhance sampling between kinetically separate regions. Therefore, these CVs should correlate with the slowest structural degrees of freedom within the system. Exclusion of these slow modes leads to hysteresis and slow convergence issues^12,14^. For example, even for the simplest test cases such as capped Alanine dipeptide, hysteresis can arise if we choose the faster ψ coordinate for enhanced sampling.

Given all of these problems with enhanced sampling algorithms, we instead aim to solve a simpler problem. What if we are given unbiased MD simulations for the baseline wild type (WT) and we wish to learn the dynamics for a closely related mutant? In computational biophysics, the mutant could correspond to a change in force field, an amino acid substitution, post-translational modifications, or even an alternative drug in the case of drug-binding simulations. We expect that these mutants likely sample a similar free energy landscape, albeit with different thermodynamics and kinetics. Could we design a better sampling scheme by transferring knowledge from the WT simulation to the mutant?

Transfer learning^15^ is a method from the machine learning literature where knowledge learned from modeling one task is transferred to the model for the purpose of learning another task. We wish to replicate a similar effect in molecular modeling where we transfer the knowledge learnt from a protein’s wild type to a simulation of its mutant. Ultimately, we aim to efficiently sample the mutant to predict affects of force field changes, post translation modifications, and/or amino-acid substitutions etc.

The idea of knowledge transfer is not new in computational biophysics. Researchers constantly use homology modeling^16^ to create models for systems which have not been crystallized or select CVs for enhanced sampling simulations^10^ based upon an intuition learnt from failed runs, literature search, or previously published modeling work on homologous systems. However, this is often done in an ad-hoc or heuristic fashion. For example, it might be difficult to find the “right” template for homology modeling when a large set of similar sequence identity structures are available.

We hypothesize an efficient use of transfer learning would maximally leverage the *reaction coordinates, thermodynamic, and structural* information contained in the WT simulation. Our *key* results stem from recognizing that protein mutants sample a similar set of free-energy minima connected via similar slow modes. Our model assumes that these slow modes involve the same set of residues across the WT and mutant sequences and all that remains are identifying those slow modes (Figure 1) in the WT simulation^12^ and transferring them on to a mutant simulation.

We propose transferring information from the WT’s tICA (time-structure based independent component analysis) model and MSM (Markov state model) to the mutant Metadynamics or Umbrella sampling simulations (Figure 1). tICA is a dimensionality reduction technique^17–20^ capable of finding reaction coordinates(tICs) within the dataset. are kinetic models of protein dynamics that model the dynamics as memory-less jump processes. tICA was initially used as a dimensionality reduction process^20^ for defining the Markov models’ state space though it was later shown that both tICA and MSM solve the same problem^21^ of approximating the underlying transfer operator, albeit with a differing choice of basis. The tICA^18–20^ method has non-linear^17^ extensions available which significantly improve its descriptive abilities. Furthermore, a variational principle^21^ for tICA and MSMs allows a researcher to systematically validate^22^ modeling parameters to potentially integrate out subjective modeling decisions. We recently showed that these tICs^20^ provided a set of excellent CVs for enhanced sampling via Metdaynamics^12^ or other schemes. Therefore, we hypothesize the answer lies in transferring these tICs over from one simulation to another.

But how do we transfer these slow tICA coordinates? At this point it is worth recalling that tICA is a linear combination of input features^12,17,18,20,23^. These input features are a set of real numbers encoding the protein’s conformational state and concretely might be dihedrals or contacts or RMSD to a set of landmark points. Furthermore, these features might be the result of a non-linear transform such as a Guassian kernel^12,23^. Therefore, what we wish to compute are these protein structural features for a new closely related sequence (Figure 1). For this, we will need to determine a set of features that can be applied to both the WT and mutant system after performing a structural or sequence alignment (Figure 1). For example, this might involve figuring out the equivalent atom indices for backbone dihedrals/contact distances/rmsds etc that make up the set of features used to construct the WT’s tICA model. Once such a mapping has been established, it is straightforward to transfer the linear combinations that make up the slowest modes for enhanced sampling simulations. In practice, we find we only have to modify small parts of input scripts that are fed into Plumed^24^ for performing the enhanced sampling simulations.

Our method explicitly makes the following set of assumptions:

1). The wild type and mutant proteins occupy similar set of configurations in phase space, are connected via similar pathways, and have a similar set of slow modes. This implicitly assumes equivalent structural features exist for both the WT and mutant proteins.

2). The wild type simulation captures a large portion this accessible phase space, and tICA and MSMs correctly enumerate these slowest modes.

There has been some previous work in using MSMs for efficient sampling of protein mutants. In particular, Voelz et al.^25^ used an information theoretic approach to find maximally surprising changes to a mutant MSM for performing new rounds of iterative sampling. However, their approach requires at least partial convergence of a rudimentary mutant MSM before such comparisons can be made. The amount of sampling required to make this rudimentary MSM could easily exceed the sampling of the WT, e.g. if the mutation slows down the dominant kinetics by an order of magnitude. Furthermore, at least initially, the rudimentary mutant MSM is likely to have large statistical uncertainties, potentially leading to false positives for the suprisal/self-information distance metric proposed in the paper^25^. Here, we are approaching the mutant problem from a fundamentally different perspective that aims to cannibalize all available data in the WT MSM.

**Figure 1:**
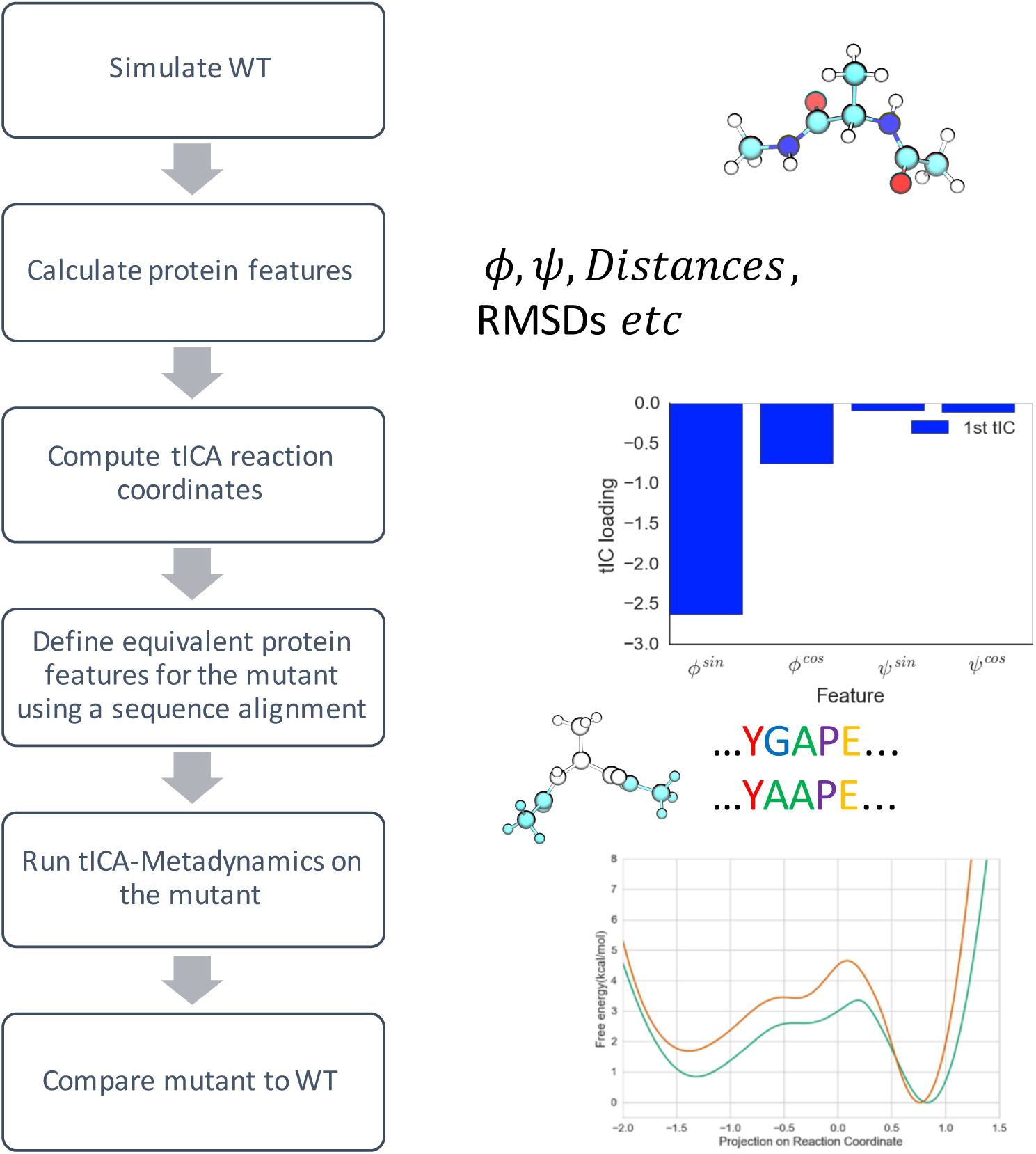
Pictorial representation for the presented method. Starting from the WT simulation, we first calculate wild type protein features. We then reduce the dimensionality keeping only the slowest modes in the system. We then transfer the tIC loadings to the mutant for faster sampling of the mutant.

### Transferable tICs are an efficient method to sample mutations

We begin by showing as a simple proof of concept that the dynamics of solvated alanine dipeptide can be re-captured across several FFs after learning the slowest modes in the “WT” model (Amber99sb-ildn^8^). We used a previously generated dataset^26^ that contained 4*μ*s of capped Alanine dipeptide run using the Amber99sb-ildn force field (FF)^8^. We then trained a tICA model on the backbone dihedrals at a lagtime of 1ns. As shown in Figure 2a, the tICA model captures the slowest mode as corresponding to movement in and out of the *α*_L_ basin while the next mode is flux in and out of the *α*_R_ basin. We next ran bias-exchange^27^ tICA-Metadynamics simulations in 3 different FFs (Amber99sbiln, Charmm27, and Amber03). The exact parameters for the well-tempered Metadynamics runs are given in SI table 1, though we empirically found that a range of parameters worked. All MD trajectories were run in the NPT ensemble with a MonteCarlo Barostat (1 atm), a Langevin integrator (300 K), and a 2 fs timestep. We used the PME method^28^ to handle long range electrostatics using a 1nm cutoff. The simulations were performed on GPUs using OpenMM^1^ and Plumed^24^. After running the Metadynamics simulation, we combined the data across the two tICs using Multi-state Bennett Acceptance Ratio (MBAR)^29,30^ algorithm. For each simulated frame, we used the last reported bias across the tIC CVs as an estimate for input into the MBAR algorithm.

The results are given in Figure 2b and 2c. We explicitly projected the Charmm27 and Amber03 datasets^26^ using Amber99sb-ildn’s state decomposition, allowing us to compare the models across force fields without having to worry about state equivalence. It can be seen that our sampling scheme efficiently learns the differences between the dynamics upon mutating the force field from Amber99sb-ildn to Charmm27 or Amber03 (Figure 2b). For example, the *α*_L_ basin in Amber03 is significantly higher in free-energy (Figure 2c) compared to Amber99sb-ildn and Charmm27.

**Figure 2:**
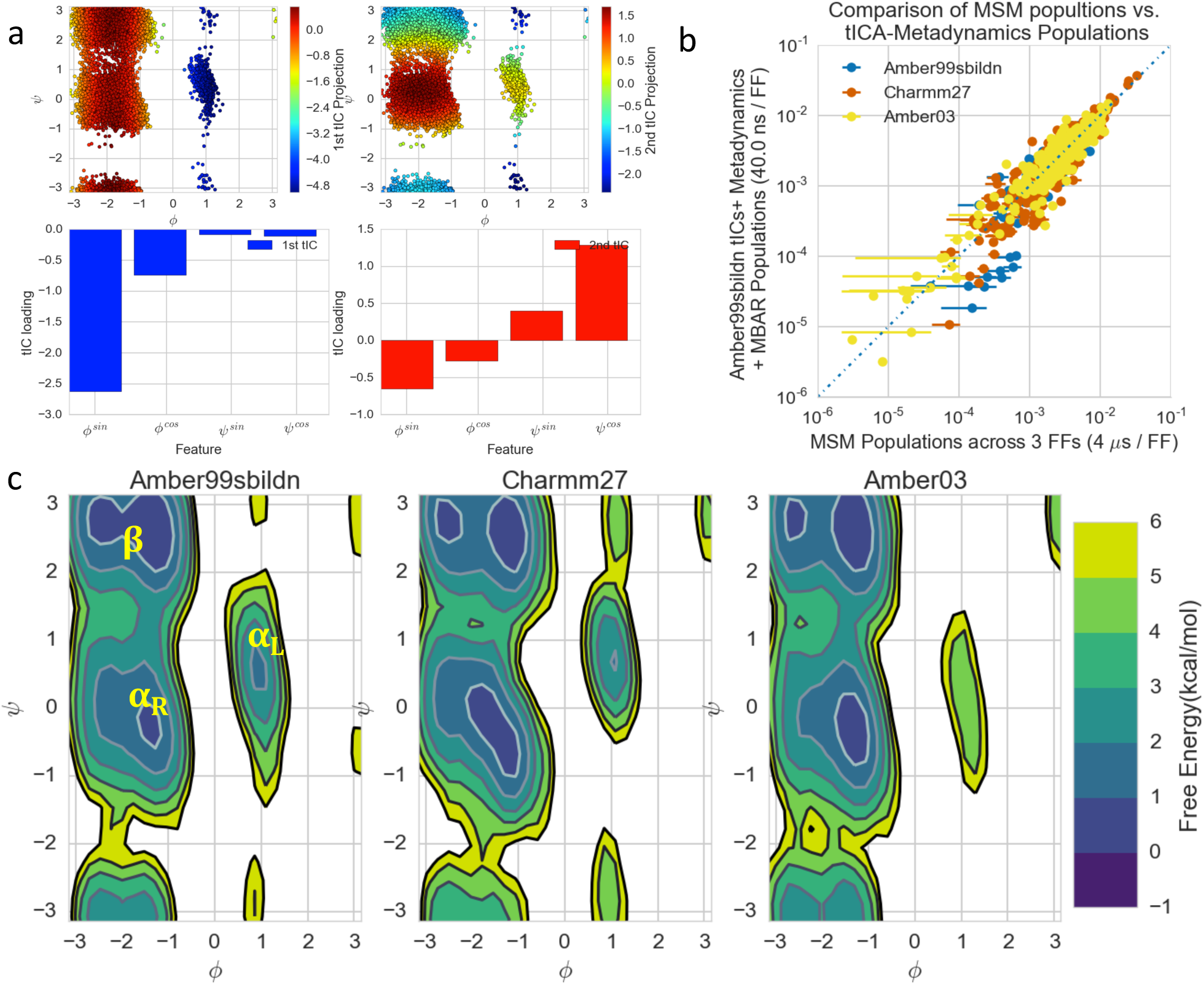
Transferable tICA-Metadynamics is an efficient method to understand the effect of mutations. a). Top two slowest tIC modes for capped Alanine learnt from unbiased MD using the Amber99sbildn model. b). Scatter plot of MSM microstate populations versus the tICA-Metadynamics inferred microstate populations. The error bars represent two standard deviations and are obtained by 10 rounds of bootstrapping. We used Amber99sbildn’s state definition across all 3 FFs for a systematic comparison. c). Reweighting the starting Ambe99sb-ildn MD dataset using tICA-Metadynamics population across 3 FFs. The 0kcal/mol is defined using Amber99sb-ildn’s most heavily populated state.

### Transferable tICs are an efficient method for sampling from more expensive force fields

But if we can transfer knowledge from one fixed charge model to another, why should we not try and predict the effects of using more expensive force fields? Force fields such as Amoeba^31^ or Drude^32^ seek to extend the descriptive capabilities of classical all-atom force fields by attempting to account for polarizable charges. However, this comes at roughly 10-100x computational cost, and it is not immediately obvious whether this cost can be justified in all modeling scenarios.

To get around this, we propose transferring information from simulations performed in cheaper force fields to more expensive simulations. This would allow us to model in polarization, or perhaps even QM effects, significantly faster than having to run those simulations from scratch. To test this, we decided to transfer knowledge from our cheaper fixed charge alanine dipeptide simulations to a simulation using the far more computationally expensive polarizable Amoeba^31^ force field. Using the same Amber tICA model as in the previous section, we sought to accelerate sampling along Alanine’s slowest mode. We used exactly the same simulation and Metadynamics parameters as in the previous section (SI table 1), *but* limited our sampling to only the dominant tIC. As a control, we also ran 5 unbiased MD simulations. The trajectories were saved every 10-20ps.

Our results are shown in Figure 3 for both the regular and enhanced sampling simulations. As it in can be seen in Figure 3a, running Metadynamics along the Amber tICs whilst using the Amoeba FF allows us to observe multiple transitions (tIC value <= -4) in and out of the *α*_L_ basin. In contrast, across the 5 control simulations with an aggregate of 17x more sampling, only a single cross over into the *α*_L_ basin (tIC value <= -4) was obtained. Such sparse regular MD statistics are likely to lead to large population uncertainties making it difficult to understand whether the effect we are seeing are due to inclusion of polarization or not. In contrast, transferring knowledge allows us to cheaply include polarization effects without sacrificing computational speed. We also note that it is perfectly possible to go in the reverse direction i.e. learn the slowest modes from the more expensive force fields and sample them using the cheaper ones.

**Figure 3:**
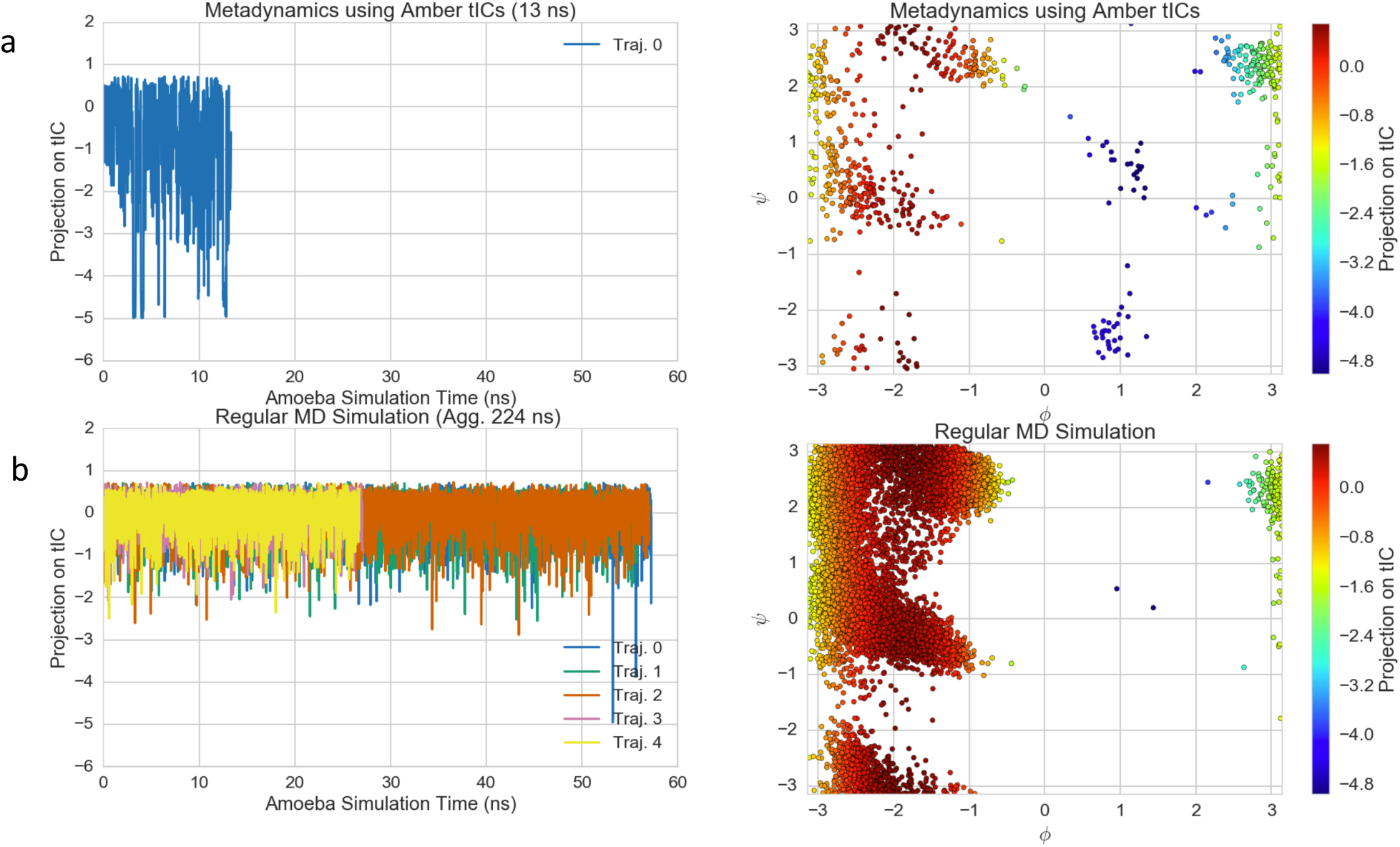
Transferrable tICs are an efficient method to include polarization effects into Alanine simulations. A). Results from running 13ns of Metadynamics using the Amoeba force field and Amber’s dominant tICA coordinate shows multiple cross-overs (tIC value <= -4) along the slowest mode. B). In contrast, across 5 trajectories with 17x more sampling, only a single crossover is obtained. The figures on the left correspond to the time traces of individual trajectories. The figures on the right are the aggregated data projected along the Ramachandran plot.

### Transferable tICA-Metadynamics can use Wild type simulation’s structural data by coupling to a MSM structural reservoir

Up to this point, our modeling efforts have only focused on using the slow tICs within the WT simulation for efficiently sampling the mutant. This might be sufficient for small peptides systems but is unlikely to work for large systems due to for example missing structural features in the construction of our tICA coordinates. While we could systematically improve the quality of our tICA model via the variational analysis^21^, there is always a finite chance of missing structural degrees of freedom. To overcome this, we recommend coupling the Metadynamics simulations to a structural reservoir containing structures sampled from the WT MSM simulation (Figure 4a). The mutant sequence is mapped onto the WT MSM states via homology modeling^16^ creating a folder of possible mutant states. Then, all that remains is creating a proposal distribution and an acceptance criterion for inserting the MSM state into the mutant Metadynamics simulation (Figure 4a). Ordinary Bias-Exchange^10,27,33^ swaps protein coordinates according the following criterion:

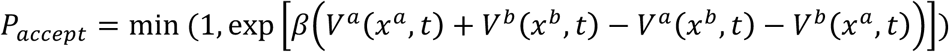

where *V^a^ (X^a^, t)* is the Metadynamics bias potential acting on coordinates, *x^a^*, of replica a at time t. However, since a MSM structural reservoir has no external bias acting on it, we change the swap probability to an insertion (from MSM to Metadynamics) probability:

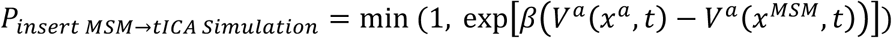

where *x^MSM^* are the coordinates for the MSM state under consideration. If accepted, the MSM state is put into the mutant Metadynamics simulation. Given enough sampling, this scheme resembles a Metropolis step. To improve the acceptance probability, we used the WT Markovian transition model to *propose* a transition state after figuring out the mutant’s current MSM state within the simulation. Using the WT transition matrix provides an excellent proposal distribution since we hypothesize that the mutant only minimally perturbs certain elements of the matrix. Our reservoir approach is similar to the high-temperature reservoir introduced by Okur et al^34^, though in this instance, the ensemble of structures is obtained via a regular MD run, and the proposal is dealt using the WT transition matrix. While the WT MSM transition matrix serves as an excellent proposal distribution it is also possible to use other proposal distributions such as the uniform distribution. Furthermore, several sampling techniques^33,35^ from the MonteCarlo literature such as the Wang-Landau scheme can be employed as well. We explicitly note that for mutant simulations, generating this MSM state reservoir would require additional steps of homology^16^ modeling, minimization and equilibration, though this is a pleasantly parallelized problem.

**Figure 4:**
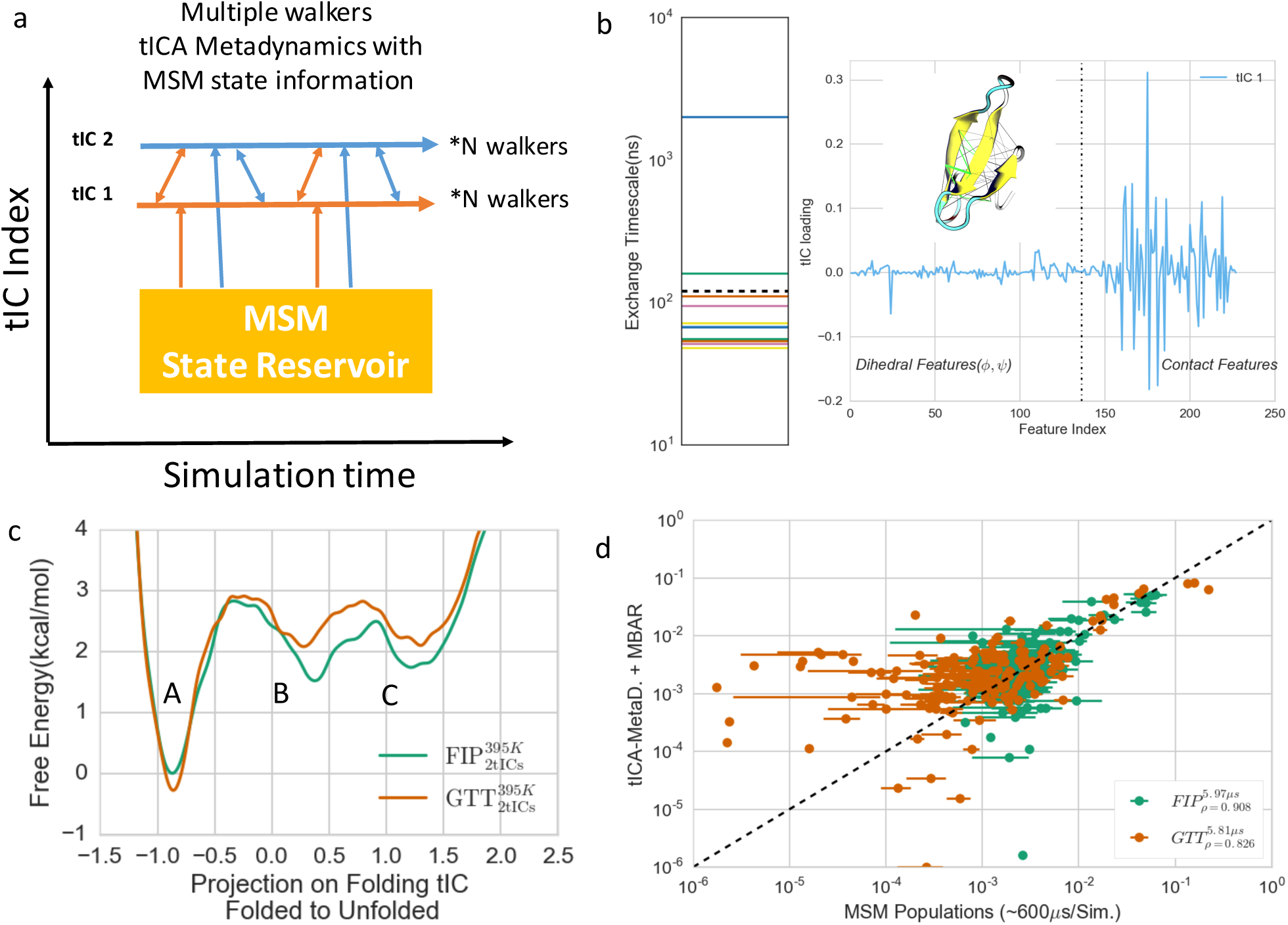
tICA-Metadynamics can be used to understand the effects of mutations upon folding simulations. a). Model of the multiple walkers’ swap scheme used within this example. The double sided arrows indicate swap attempts while the single sided arrows indicate insertion attempts from the MSM state reservoir. For mutants, homology modeling and equilibration will be necessary to generate the pre-equilibrated MSM state reservoir. b). Left Panel: Exchange timescales for the backbone dihedral and contact tICA model built using the Anton WW-FIP mutant simulation data. We decided to sample every coordinate up to the second one (dashed line) because all other modes had exchange timescales <= 100ns. b). Right Panel: tIC loading for the dominant folding tIC shows that it is a complex combination of both the WW’s backbone dihedrals and contacts. The inset shows the contact distances used to build the model with the most important (highest tIC loading) distances shown in green. c). PMFs obtained from the Metadynamics simulations using the scheme highlighted in a). The GTT mutants’ unfolded state is de-stabilized relative to the folded state. In both cases, we used the WW-FIP mutant to define the 0kcal/mol. The total amount of sampling across all the walkers was about 5-6μs per mutant. d). Comparison of the Anton-MSM state-populations vs. the MBAR reweighted tICA-Metadynamics results. The MSM error bars represent two standard deviations obtained via 10 rounds of bootstrapping. The legend shows both the sampling per mutant, and the correlation coefficients between the MSM thermodynamics vs the Metadynamics populations. Protein images were generated using VMD^36^, while the graphs were generated using IPython Notebook^37^ and MSMExplorer^38^.

Lastly, it is possible to use a neutral replica^27^ within this setup. The neutral replica has no external bias acting on it and approximately samples from the canonical distribution. However, if a neutral replica is used, we recommend only allowing the neutral replica to swap with the biased replicas since an appropriate asymptotically correct swapping criterion for swapping between the neutral replica and the MSM state reservoir doesn’t exist.

We tested our methodology by predicting the effects of the GTT mutation upon the folding of the WW domain^39,40^. We began by learning a tICA model (50ns lagtime) on the backbone dihedrals and selected contacts for the WT mutant (WW-FIP). We kept the top 15 tICs, and made a MSM at a 50ns lag time on a 200 state model. Our tICA model indicated that the slowest mode (Exchange time scale > 1 *μs*) corresponded to the folding while the second slowest mode (Exchange time scale > 100 ns) corresponded to formation of an off-pathway register shifted state (Figure 4b-c). Since every subsequent slow tIC mode has exchange timescales of less than ∼100ns, we chose to focus our sampling on these two tICs. We ran the simulations for *both* the FIP35 WT protein and the GTT triple mutant for a more systematic comparison. Similar to previous work^39^, all the simulations were performed in the NVT ensemble with a 2fs time step at 395K. We used the PME method^28^ to handle long range electrostatics using a 1nm cutoff. The simulations were performed on GPUs using OpenMM^1^ and Plumed^24^. After running the Metadynamics simulations, we used MBAR to re-weight to the MSM state space and obtained the PMFs along the dominant tIC. All relevant simulations parameters and reweighting details^41,42^ are shown in SI Table 2 and SI Note 1.

The results for both of our enhanced sampling simulations are given in Figure 4c-d. Two different insights emerge from our enhanced sampling scheme relative to the Anton^39^ results (SI Figure 1). Similar to the Anton simulations, our FIP unfolded state (Figure 4d, tiC value >-0.25) has a distinct two state behavior. Basin ‘C’ corresponds to the unfolded and collapsed state. This basin also includes an off-pathway register shifted state. The second high free-energy basin (Figure 4d, B) is an on-pathway intermediate state where two of the three beta-strands have formed. The unfolded state in our ensemble is more populated than in the Anton simulations. These on and off-pathway intermediate states were not detected in the original two-state folding reaction coordinate for the WW domain^39,40^ though it was later found from the simulations using a variety of techniques^43^. We note that our tICA analysis was able to identify the on-pathway folding intermediate and the off-pathway state as the top two slowest modes (tICs) within our model.

As can be seen in Figure 4c-d, our simulations indicate that the GTT mutant de-stabilizes the unfolded state and the on-path intermediate state, leading to increased folded population. According to our calculations, the population for the folded state (defined as dominant tiC value <=-0.25) increases from 55% to 67% of the ensemble when going from FIP to GTT indicating a ∼0.4kcal/mol stabilization. These results are comparable to the ∼0.65 kcal/mol stabilization (58% to 76%) for the unbiased Anton MD trajectories. These results are also in line with previous experimental work^39^ though our simulations required about 100-125x less aggregate sampling(∼5-6*μs* vs 600*μs*). More importantly, the current sampling was performed in parallel so that no single walker had to be run for more than 50-200ns (∼3-7 days on K40 GPUs using OpenMM^1^ and Plumed^24^). We also believe it might be possible to optimize this further by modifying the Metadynamics parameters and Metropolis swap schemes/rate.

### Transferable tICA-Metadynamics can use Wild type simulation’s thermodynamic data as a prior for the underlying free energy landscape

Lastly, we turn to efficiently using the thermodynamic information contained in the WT simulation. To that end, we recommend using the WT simulation to identify minimum values along each tIC coordinate, aka the thermodynamic minima, to plug into a variant of Metadynamics, namely Transition-Tempered Metadynamics (TTMetaD)^44^. In TTMetaD, the Gaussians heights are scaled according to the number of trips between basins. We also believe that it is possible use to the WT free energy surface as a Bayesian^45^ prior for the mutant Metadynamics simulation, though that is beyond the scope of this work. The latter might involve starting off with a ‘partially’ constructed free energy-landscape such that the Metadynamics engine only has to fill in the regions that are different between the WT and the mutant.

## Discussion and Conclusion

Our current results open up several interesting avenues for future work. For example, up to this point, we have only focused on enumerating the thermodynamic differences between the mutants. However, the recent work in kinetic reweighting either via Maximum caliber^46^, TRAM^47^, or plain transition state theory could potentially be used to obtain the an estimate for the mutants’ perturbed kinetics. This raises the intriguing possibility of getting estimates for both the kinetic and thermodynamics of a mutant simulation for a miniscule fraction of the WT’s compute cost. An excellent application for this would be the ability to predict changes in a drug’s binding and unbinding kinetics. Our approach explicitly includes all of the protein’s slow conformational modes, in addition to the drug binding mode—making it more accurate.

One possible problem with our current approach is the determination of how far we can move away from the WT in sequence space before the transfer approach fails. Are the tICs learnt from a WT simulation applicable to a sequence with minimally sequence similarity? What is the distance metric and how do we define minimal? A similar problem is faced in homology modeling, where the quality of the model depends on the underlying sequence conservation. It is possible that the heuristic value of 40-50% sequence identity cutoff used in homology modeling might be applicable here too, but we concede that that value is simple conjecture at this point.

A more involved solution to this problem is to consider clustering the entire sequence super family. For example, there are 518 known human kinases^5^. One could potentially cluster the sequences using evolutionary distance metrics into *m* representative sequence clusters, where *m* is the number of possible unbiased simulations that can be performed. Those *m*-simulations are then run and analyzed via tICA and Markov models. It is worth noting that the simulations for the *m* sequences are perfectly parallelizable, allowing for synergistic collaborations between different research institutes. For all other sequences, we can then use the tICs from its closest cluster center or perhaps even combine the tICs from the k nearest neighbors.

To summarize, we present a new method Transferable tICA–Metadynamics for the efficient sampling of protein mutations by transferring the reaction coordinates, structural, and thermodynamic data from the WT simulation to the mutant. Our method explicitly assumes that the WT and the mutant share a similar set of slow modes. Under this assumption, we then show that the slow modes of the WT can be transferred to the mutant simulation by computing an equivalent set of protein structural features. This requires using a protein structural alignment to identify equivalent residues which is readily possible using modern software^48,49^. We benchmarked our method on two test cases showing how switching force field in alanine dipeptide causes shifts in the propensity and location of the *α*_L_ basin, and recapturing the previous results that the GTT mutant of WW domain stabilizes the active state.

## Acknowledgements

The authors would like to thank various members of the Pande lab for useful discussions and feedback on the manuscript. M.M.S would like to acknowledge the authors from^41^ for their help regarding re-weighting of the Metadynamics simulations. M.M.S would like to acknowledge support from the National Science Foundation grant NSF-MCB-0954714. This work used the XStream computational resource, supported by the National Science Foundation Major Research Instrumentation program (ACI-1429830).

## Supporting Information Available

The SI contains the Metadynamics simulation parameters, projection of the FIP WT simulation on to its dominant reaction coordinate, and a note about re-weighting.

## Code and data availability

All the code needed to reproduce the main results of this paper is available at https://github.com/msultan/tica_metadynamics.

## TOC Graphic

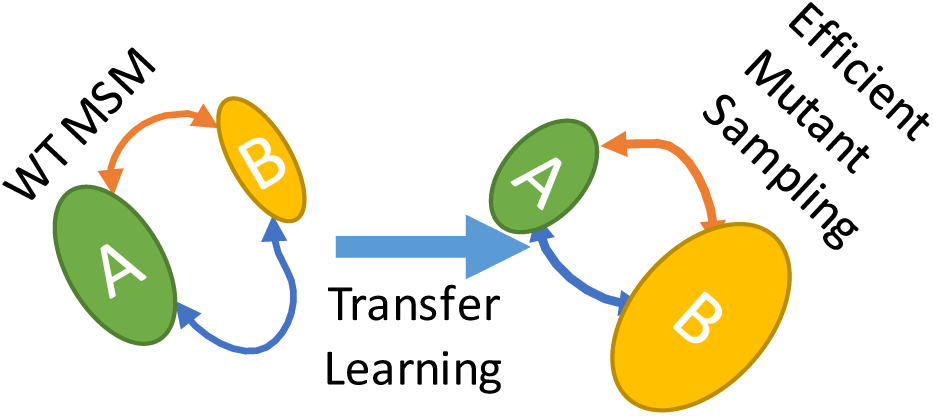

